# Temporary suspension of mineral phosphorus reduces mobilizable bone zinc in adult laying hens irrespective of the dietary zinc supply

**DOI:** 10.1101/2024.12.28.630579

**Authors:** Annegret Lucke, Agnes Weller, Julia Pokorny, Reinhard Puntigam, Julia Slama, Karl Schedle, Wilhelm Windisch, Daniel Brugger

**Affiliations:** Institute of Animal Nutrition and Dietetics, Vetsuisse-Faculty, University of Zurich, Winterthurerstrasse 270, 8057 Zurich, Switzerland; Chair of Animal Nutrition, TUM School of Life Sciences, Technical University of Munich, Liesel-Beckmann-Strasse 2, 85354 Freising, Germany; Institute of Animal Nutrition, Livestock Products, and Nutrition Physiology, Department of Agrobiotechnology, University of Natural Resources and Life Sciences, Muthgasse 11, 1190 Vienna, Austria; South Westphalia University of Applied Sciences, Lübecker Ring 2, 59495 Soest, Germany; Animal Nutrition and Nutritional Physiology, Faculty of Agricultural and Environmental Sciences, University of Rostock, Justus-von-Liebig-Weg 6b, 18057 Rostock, Germany

## Abstract

This study examined the effects of short-term dietary zinc (Zn) and phosphorus (P) variations on the mobilizable bone Zn pool and overall Zn status in adult laying hens. Forty-eight hens (50% Lohmann Brown Classic, 50% Lohmann LSL Classic) were housed in pairs (one hen per breed per pen) across 24 pens. The pens were randomly assigned to one of two dietary P levels (0.37% or 0.84% in DM) using a high-protein corn-soybean diet (11.4 MJ AME/kg, 21.5% CP) during a 14-day acclimatization period. Following acclimatization, pens from both P groups were further randomized into four dietary treatments in a 2 × 2 factorial design, varying in P levels (low vs. high) and Zn supplementation (28 vs. 131 mg/kg) over an 8-day experimental feeding phase. Performance metrics, egg production and quality, and tissue mineral concentrations (plasma, liver, bone, and eggs) were measured. Statistical analyses were performed using linear mixed models in SAS 9.4, incorporating random effects of pen nested within treatment group and fixed effects of dietary P, dietary Zn, breed, and their interactions. Tukey-corrected 95% confidence intervals were used to estimate effect differences, with significance set at P < 0.05. Performance metrics, including egg production and body weight, were unaffected by dietary treatments (P > 0.1), indicating no clinical symptoms of Zn deficiency. However, hens on low-Zn diets exhibited significant reductions in plasma Zn concentration (-0.83 mg/L; P = 0.0008) and liver Zn concentration (-6.78 mg/kg DM; P = 0.01), confirming subclinical Zn deficiency. Low-Zn diets also increased the femoral molar Ca:P ratio by 0.15 (P = 0.01), irrespective of dietary P supply. Interestingly, low-P diets led to a significant reduction in femur Zn content (-0.46 mg; P = 0.0009), regardless of Zn supplementation, following 21 days of reduced P feeding. These findings highlight the higher susceptibility of laying hens to phytate antagonism compared to broilers, as evidenced by measurable subclinical Zn deficiency under short-term Zn deprivation. Additionally, a temporary suspension of mineral P supply appeared to impair the mobilizable bone Zn pool. The underlying functional mechanisms driving these interactions remain unclear and warrant further investigation.

## Introduction

Organisms exhibit complex regulatory mechanisms to maintain zinc (Zn) homeostasis. Research across different species has shown the complexities of zinc (Zn) metabolism. For example, rats adjust their Zn metabolism in response to varying dietary Zn levels, by decreasing half-life time of Zn exchange while maintaining a constant mobile Zn pool, illustrating the ability to modulate Zn exchange kinetics effectively (Windisch and Kirchgessner, 1995). Brugger et al. (2014) identified that the dose-response patterns of apparently digested amount of Zn and liver Zn levels to gradual variation in diet Zn serve as effective markers for identifying the transition from deficient to sufficient Zn supply in a piglet model of short-term subclinical Zn deficiency, which has been validated in follow-up studies (Boerboom et al., 2022). Comparable response patterns have been observed also in broiler chickens and cattle (Schwarz and Kirchgessner, 1975; Boerboom et al., 2021).

In laying hens, Paulicks and Kirchgessner (1994) found decreased feed intake and egg production as well as a drop in whole egg Zn while feeding a long-term (48 weeks) Zn deficient diet, emphasizing Zn’s critical role in poultry productivity. Despite these findings, knowledge regarding Zn dynamics and requirements in laying hens (and broilers) remains limited, particularly concerning the effects of short-term fluctuations in Zn supply and the impact of dietary phytate on Zn bioavailability. To this day, many studies have been published on the efficacy of different Zn sources in poultry feeding. Most of these papers discuss the findings by extrapolating lessons learnt from monogastric mammals to the birds. However, since not much is known on chickens’ quantitative Zn metabolism and its functional background, these conclusions are in best case speculative. Research in poultry zinc metabolism is hampered by the absence of standardized and reproducible experimental models to study this trait, like they have been developed thus far for rats and pigs (Windisch, 2003; Brugger et al., 2014).

Before developing experimental models to precisely study Zn requirements in laying hens, it is essential to consider their unique metabolic demands (e.g. during egg production). Phytate, the primary storage form of phosphorus in plant-based feedstuffs, is well-documented to chelate divalent cations like Zn, thereby reducing its absorbability by forming insoluble complexes in the gastrointestinal tract (Humer et al., 2015). Broilers have been shown to efficiently degrade phytate, primarily due to a combination of endogenous and microbial phytase activity, and this digestive capacity is reduced when feeding mineral P sources like mono-Ca-phosphate (MCP) (Zeller et al., 2015; Ingelmann et al., 2019; Ma et al., 2019; Sommerfeld et al., 2019). Compared to broiler chickens, phytate P is less available in laying hens, which may be related to higher amounts of Ca in layer diets (Ma et al., 2019). The enhanced phytate degradation may allow broilers to absorb zinc more effectively, mitigating the negative effects of phytate on Zn bioavailability and explaining observed differences to weaned piglets, which apparently are much more susceptible to the phytate antagonism (Schlegel et al., 2013). Moreover, the high Ca content in layer diets may exacerbate complexion of phytate with Ca and Zn, thus decreasing Zn availability (Humer et al. 2014). Consequently, while broilers can in part compensate for phytate’s inhibitory effects, laying hens may be more susceptible to reduced Zn absorption, necessitating careful dietary formulation to ensure adequate zinc levels for optimal health and productivity in egg production. Furthermore, understanding the temporal dynamics of zinc metabolism in response to short-term variations of Zn supply is important to develop a robust model that accurately reflects the Zn requirements of laying hens, addressing their specific physiological requirements (Brugger et al., 2022). Previous research in piglets successfully established a model for subclinical zinc deficiency, which included a 14-day acclimatization phase with a diet formulated to meet their nutritional requirements, followed by an 8-day period of varying zinc supply in a phytate-rich diet (Brugger et al., 2014; Boerboom et al., 2022). Similar models are the basis to investigate metabolic mechanisms without confounding influences of clinically manifest Zn deficiency (Brugger and Windisch, 2017). While previous studies indicate that four months are required to induce clinical zinc deficiency in laying hens (Paulicks and Kirchgessner, 1994), models to assess short-term subclinical zinc deficiency, which is supposed to be the more common Zn deficiency phenotype in humans and animals (Nielsen, 2012), remain to be developed. Furthermore, the work by Paulicks and Kirchgessner (1994) does not reflect the current layer genotypes and it could be argued that today’s high-performing breeds might be more susceptible to fluctuations in dietary Zn supply.

Based on the aforementioned reports in broiler chickens, according to which the mineral P supply in the diet affects phytate breakdown in the gastrointestinal tract, this pilot study was designed to pave the way towards the development of a subclinical Zn deficiency model in laying hens. Our working hypothesis was that chickens fed without mineral phosphorus for 21 d will have a higher endogenous Zn status (bone Zn, liver Zn, plasma Zn) than a control with MCP in the diet, based on the assessment of Zn in liver, plasma, and bone. Furthermore, we additionally varied the diet Zn supply (analysed: 28 vs. 131 mg/kg) for the last 8 d to further understand the interaction between P and Zn metabolism to test the susceptibility of layers to a short-term suspension of diet Zn supplementation. The timeframe of 8 d for the native Zn challenge mimicked our earlier attempts in pigs. Since piglets are more susceptible to diet induced Zn deficiency than (broiler) chickens (Schlegel et al., 2013) we assumed that pathophysiological events would therefore be unlikely and any reduction in body Zn status would be of a subclinical nature. Since earlier studies suggest the digestive capacity of layer chickens for phytate differs between breeds, this study combined adult Lohmann Brown Classic and Lohmann LSL Classic hens. The results from this study have the potential to guide future research on layer Zn metabolism and feeding and to pave the way towards the development of a subclinical Zn deficiency model in layers.

## Material and Methods

This animal study was reviewed and approved by the responsible animal welfare officer and was registered and approved with the responsible Austrian authorities (BAES-FMT-FV-2018-008). No animal showed any sign of pathology, nutrition-associated or otherwise, during the whole study period.

### Animals, diets, and experimental design

Forty-eight adult laying hens (50% Lohmann Brown Classic, 50% Lohmann LSL Classic, average life weight 2109 ±220 g and 1796 ±84.2 g, respectively) were obtained from a commercial producer. Animals from both breeds were randomly assigned over 24 wooden pens (1.5 m^2^) with spelled husks on concrete floor as bedding material. Each pen contained one hen from each breed (n = 2 hens pen^-1^). Pens were equipped with nipple drinkers and troughs, allowing pen-wise determination of feed intake as well as perches and nest-boxes.

Pens were randomly assigned to one of two groups, receiving 14 d of acclimatization feeding with a basal diet (11.4 MJ AME/kg, 21.5% CP) based on corn, soybean meal (44% CP), and potato protein (Table 1). This basal diet did not contain exogenous phytase or crystalline amino acids, and met or exceeded recommended nutrient levels (Lohmann, 2020a, 2020b) including Zn (added Zn: 100 mg/kg from analytical grade C₄H₆O₄Zn * 2H₂0; analyzed Zn: 128 mg/kg) but not phosphorus, which was varied between groups (added P: 0% or 0.52%; analyzed P: 0.37% or 0.83 %).

**Table 1.**
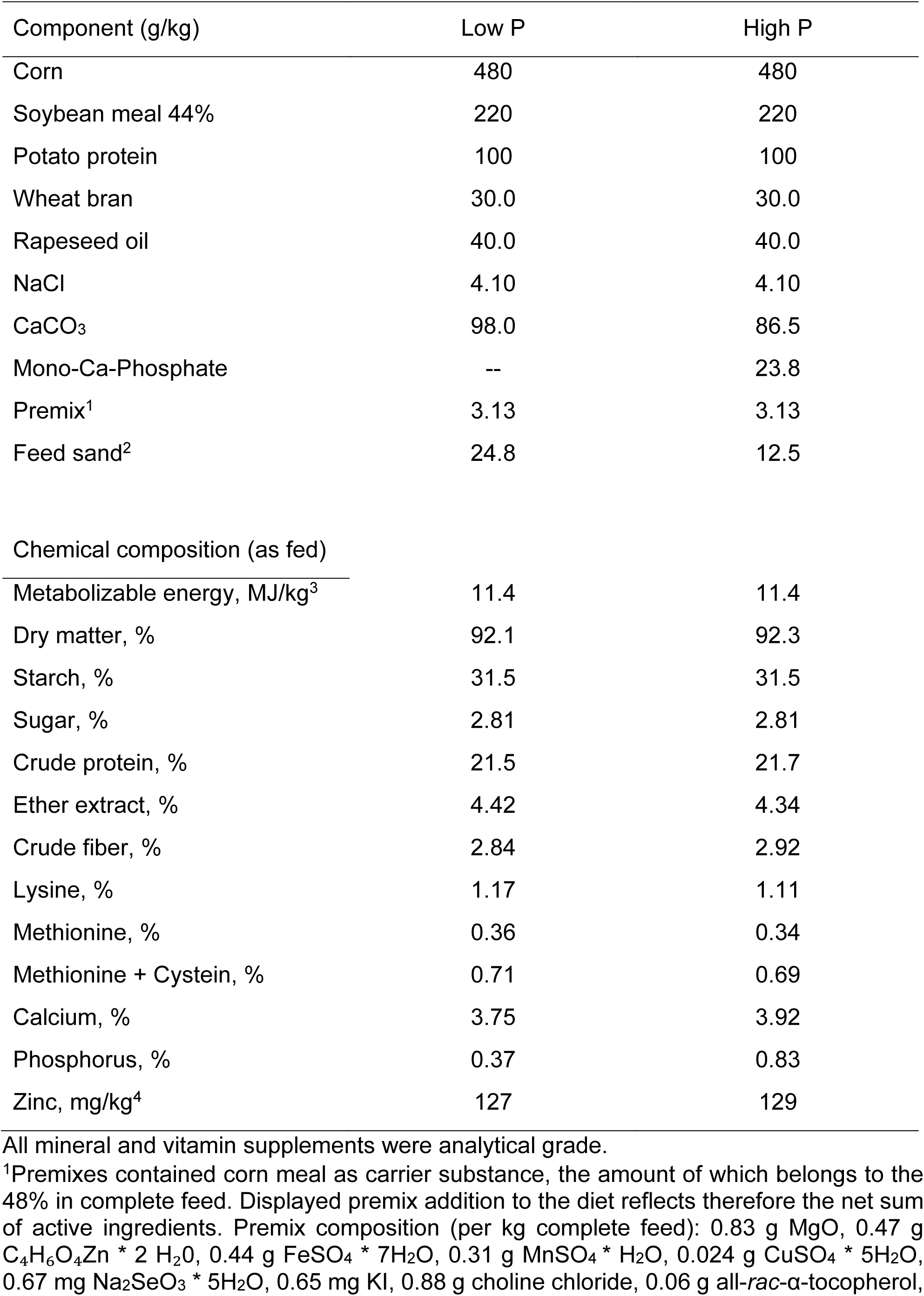
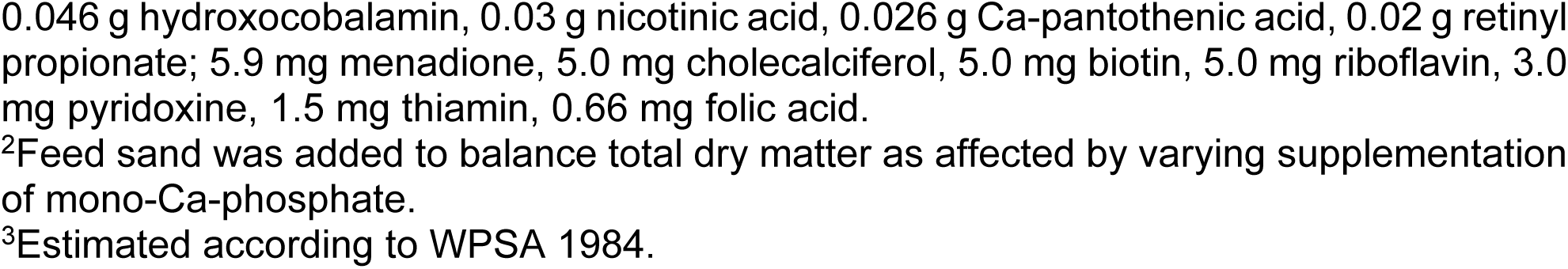
Ingredients and chemical composition of basal layer diets fed during the pre-experimental phase.

| Component (g/kg) | Low P | High P |
| --- | --- | --- |
| Corn | 480 | 480 |
| Soybean meal 44% | 220 | 220 |
| Potato protein | 100 | 100 |
| Wheat bran | 30.0 | 30.0 |
| Rapeseed oil | 40.0 | 40.0 |
| NaCl | 4.10 | 4.10 |
| CaCO<sub>3</sub> | 98.0 | 86.5 |
| Mono-Ca-Phosphate | -- | 23.8 |
| Premix<sup>1</sup> | 3.13 | 3.13 |
| Feed sand<sup>2</sup> | 24.8 | 12.5 |
| Chemical composition (as fed) |  |  |
| Metabolizable energy, MJ/kg<sup>3</sup> | 11.4 | 11.4 |
| Dry matter, % | 92.1 | 92.3 |
| Starch, % | 31.5 | 31.5 |
| Sugar, % | 2.81 | 2.81 |
| Crude protein, % | 21.5 | 21.7 |
| Ether extract, % | 4.42 | 4.34 |
| Crude fiber, % | 2.84 | 2.92 |
| Lysine, % | 1.17 | 1.11 |
| Methionine, % | 0.36 | 0.34 |
| Methionine + Cystein, % | 0.71 | 0.69 |
| Calcium, % | 3.75 | 3.92 |
| Phosphorus, % | 0.37 | 0.83 |
| Zinc, mg/kg<sup>4</sup> | 127 | 129 |

All mineral and vitamin supplements were analytical grade.
<sup>1</sup>Premixes contained corn meal as carrier substance, the amount of which belongs to the 48% in complete feed. Displayed premix addition to the diet reflects therefore the net sum of active ingredients. Premix composition (per kg complete feed): 0.83 g MgO, 0.47 g C<sub>4</sub>H<sub>6</sub>O<sub>4</sub>Zn \* 2 H<sub>2</sub>O, 0.44 g FeSO<sub>4</sub> \* 7H<sub>2</sub>O, 0.31 g MnSO<sub>4</sub> \* H<sub>2</sub>O, 0.024 g CuSO<sub>4</sub> \* 5H<sub>2</sub>O, 0.67 mg Na<sub>2</sub>SeO<sub>3</sub> \* 5H<sub>2</sub>O, 0.65 mg KI, 0.88 g choline chloride, 0.06 g all-*rac*- $\alpha$ -tocopherol,
0.046 g hydroxocobalamin, 0.03 g nicotinic acid, 0.026 g Ca-pantothenic acid, 0.02 g retinyl propionate; 5.9 mg menadione, 5.0 mg cholecalciferol, 5.0 mg biotin, 5.0 mg riboflavin, 3.0 mg pyridoxine, 1.5 mg thiamin, 0.66 mg folic acid.
<sup>2</sup>Feed sand was added to balance total dry matter as affected by varying supplementation of mono-Ca-phosphate.
<sup>3</sup>Estimated according to WPSA 1984.

After the 14-d acclimatization feeding, pens within the low P and high P (LP, HP) groups were each randomly assigned to one of two subgroups, receiving the same diet as during acclimatization or a diet with the same P level as before but without Zn addition. This resulted in four experimental feeding groups: LPLZ, LPHZ, HPLZ, and HPHZ, receiving analyzed dietary P at 0.36, 0.38, 0.87 and 0.82 % and dietary Zn at 27.3, 132, 28.6 and 130 mg/kg, respectively (Table 2). Differences in analyzed dietary P and Zn between diets receiving the same level of mineral addition were only numerical according to ANOVA validation (diet*batch). Therefore, in the further course of this manuscript, we use the average analyzed values of 0.37% and 0.84% as well as 28 and 131 mg/kg, to describe the LP, HP, LZ, and HZ levels in diets, respectively. Experimental feeding lasted for eight consecutive days after the 14d acclimatization phase, in the end of which the experiment was terminated with the slaughtering of the hens by trained personnel.

**Table 2.**
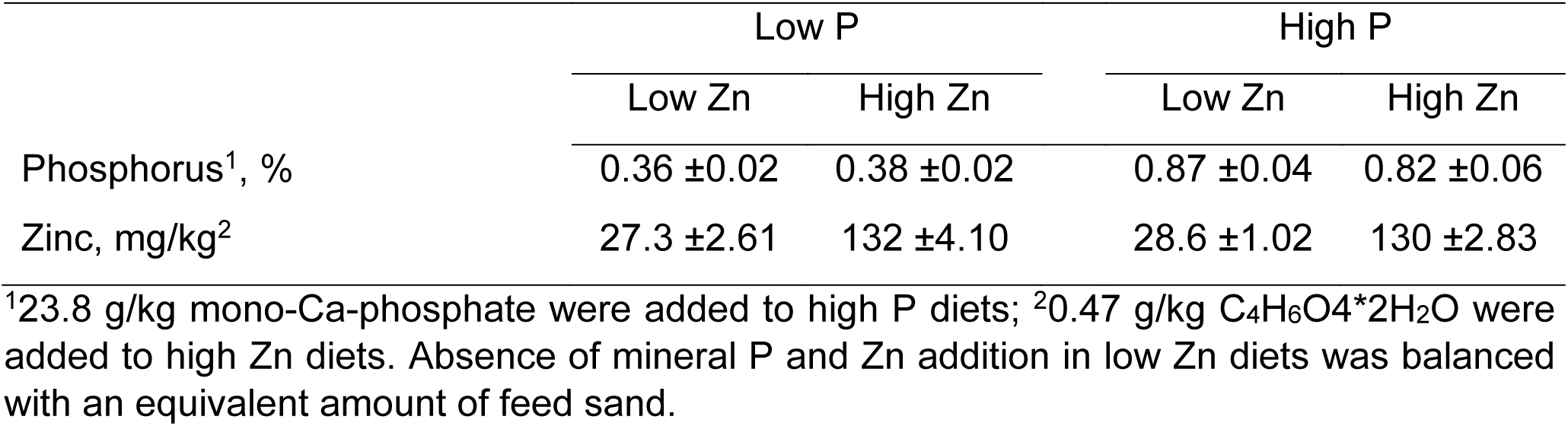
Dietary phosphorus and zinc concentrations in experimental diets (92% DM), as affected by varying addition of CaCO_3_, mono-Ca-phosphate and C₄H₆O₄Zn * 2H₂0 supplements.

### Sampling

Feed samples were stored at 5°C until further usage. During the study, feed intake was determined pen-wise per week. Hens were weighed before the onset of the study and subsequently on a weekly basis, always at 8 a.m.. Eggs were collected daily and counted for the determination of the weekly egg yield and subject to the determination of egg quality parameters.

Hens were slaughtered after eight experimental days and samples of whole blood in Li-Heparin Tubes and body tissues (complete liver, left femur) were taken. Blood samples were centrifuged (1100 x g, 10 min, 4°C) to collect blood plasma, which was stored at -20°C until further usage. Bone and liver samples were vacuum sealed in plastic bags and stored at -20°C until further usage.

### Dry matter and nutrient analyses

Diets were analyzed according to the following standard procedures (element analysis was described in detail further below): dry matter (DM, Method 3.1), starch, sugar, crude protein (CP, Method 4.1.1), ether extract (EE, Method 5.1.2), crude fiber (CF, Method 6.1.1) and amino acids (lysince, methionine, cysteine, Method 4.11.1) (VDLUFA, 2012) and the data was used to estimate metabolizable energy in complete feed according to (WPSA, 1984)

Total dry weight of whole egg was assessed on the last day of acclimatization (d_0_) and experiment (d_8_), respectively. Total livers and each left femur were taken at the time of slaughter. Livers were dried at 50°C for 2 d. After bone strength measurement (procedures shown below), the femur was dried in an oven (105°C, 1d) and ashed in a furnace (470 °C, 2d) in platinum dishes to determine the total ash weight.

### Egg weight and egg shell quality

Egg weight and egg shell quality were determined daily from the beginning of the acclimatization to the end of the experiment.

Egg weight was measured with a precision scale (440-43N; Kern) and egg shell strength with the Textur Analyzer TA.HD. plus system (Stable Micro Systems) by applying pressure (up to 750 kg) from tip to bottom. Eggshell thickness was measured with an electronic caliper (IP65; Mitutoyo) at the tip, middle part and bottom of each egg to determine an average value.

### Bone size and breaking strength

The left femur of each hen was measured for length and girth (mid of shaft) with a manual caliper. Subsequently the bone was subject to a three-point-pressure test with a 50 kg measuring cell and a 5 kg weight to determine the breaking strength of each bone with the Textur Analyzer XT Plus system (Stable Micro Systems).

### Element analyses

Feed, plasma, dried liver and bone ash (left femur, 470°C, 48 h) were subject to microwave wet digestion with 65% HNO_3_ and 10% H_2_O_2_ with the equipment and procedure described in detail by Brugger et al. 2014. The solutions were used for the measurement of Zn, P, and Ca as described below.

Calcium and Zn in feed, plasma, bone, and liver extracts was measured with ammonia as reaction gas in dynamic reaction cell mode or helium as collision gas in kinetic energy discrimination mode via inductively-coupled plasma mass spectrometry (Nexion 450D, Perkin Elmer). Zinc and Ca were measured on isotopes ^66^Zn and ^43^Ca, respectively, with ^72^Ge as external standard. For this purpose, mass spectrometry-certified single-element standard materials (N9303758, N9303733, N9303739; Perkin Elmer) were applied over the following concentration gradient for Ca, Zn, and Ge, respectively: 0.005, 0.01, 0.05, 0.1, 0.5, 1.0 mg/L.

Phosphorus in feed, plasma, bone, and liver extracts was measured with vanadate-molybdate reagent (108498, Merck Millipore) on a UV MC^2^ spectrophotometer (Safas Monaco) at 430 nm wavelength according to VDLUFA 2012.

### Statistical analyses

Data analysis was conducted with SAS 9.4 (SAS Institute Inc.) and independently for the acclimatization and experimental period. In any case, linear mixed models were estimated with the procedure MIXED including the random effect of the respective pen nested under the respective feeding group and the subsequently described fixed effects.

Feed intake per pen and week as well as for the average pen-wise intake during the total acclimatization period was analyzed for the fixed effect of “diet P”. The average bird-wise live weight, egg yield, egg weight, egg shell strength, and egg shell thickness during single weeks and the total acclimatization period was analyzed for the fixed effects of “diet P”, “breed”, and interaction.

All obtained data from the experimental period were analyzed for the fixed effects of “diet P”, “diet Zn”, “breed”, and their respective interaction, except for the average pen-wise feed intake for which the “breed” effect could not be estimated.

The estimation of effect sizes and minimum number of replicates for the present study, to reach a minimum power of 1 - β = 0.8 and α = 0.05, occurred with R-4.4.2 (The R Project for Statistical Computing) with the SIMR package (Green and MacLeod, 2016)applying earlier published data from Zn metabolic trials in laying hens (Paulicks and Kirchgessner, 1994) and weaned piglets (Brugger et al., 2014).

## Results

No animal showed clinical signs of mineral deficiency at any time of the study. In general, no pathologies were observed and animals performed as expected given their production stage.

All results tables present the least square mean values (LSmean) estimated with the respective linear mixed models including the Tukey-corrected 95% confidence limits of the respective LSmeans. Furthermore, effect differences (ΔLSmean) for the respective effects of diet P, diet Zn, and breed including the Tukey-corrected 95% confidence limits of these estimates are given.

### Diet homogeneity

The homogeneity of the diets during acclimatization and experiment was sufficient, based on the analyzed concentrations of the nutrients under study as affected by varying addition to complete feed (Table 2).

### Performance

Table 3 highlights the response of pen-wise feed intake, egg yield, and egg weight during the acclimatization and experimental phase. In addition, Supplementary Table 1 highlights the response of live weight over time.

**Table 3.**
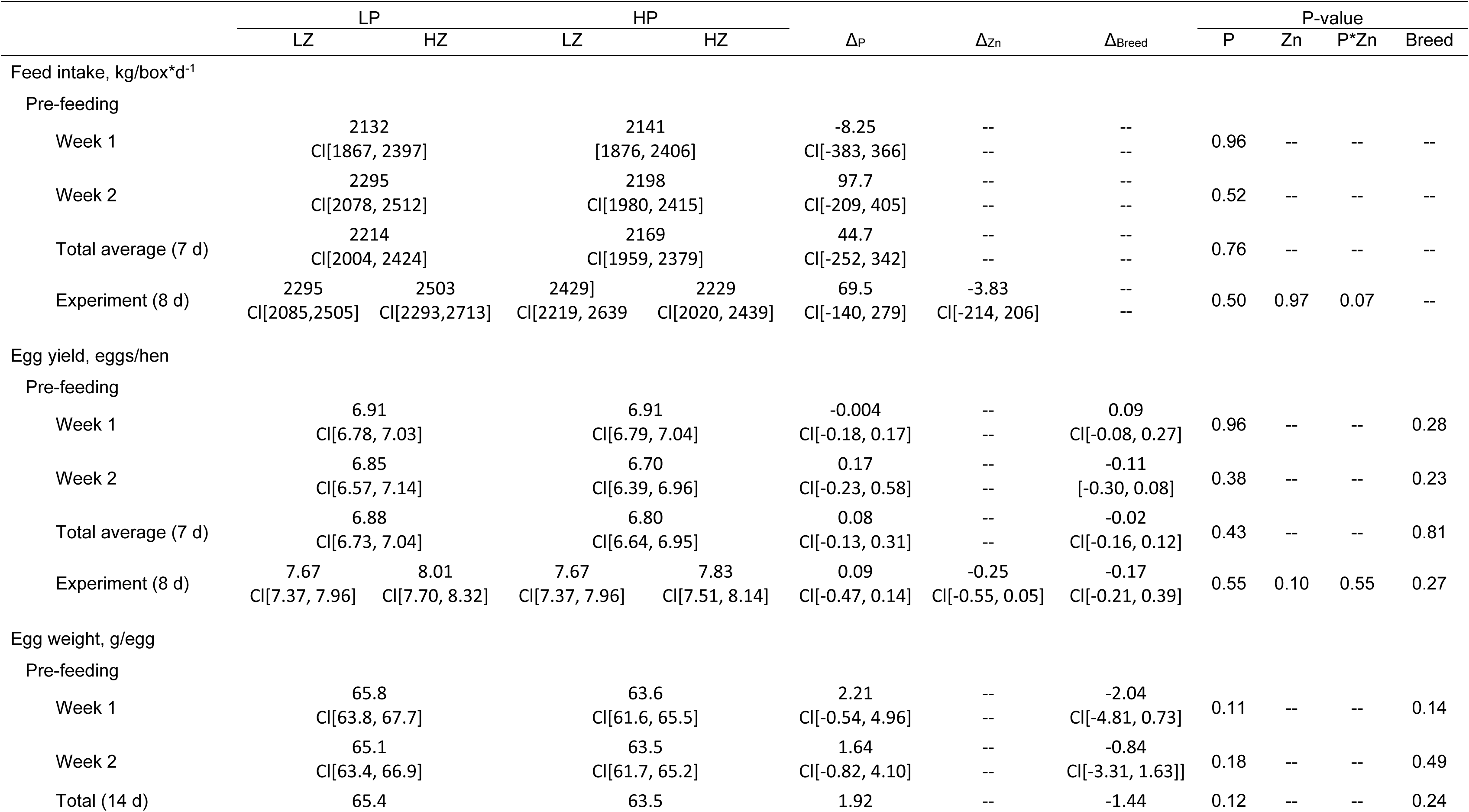
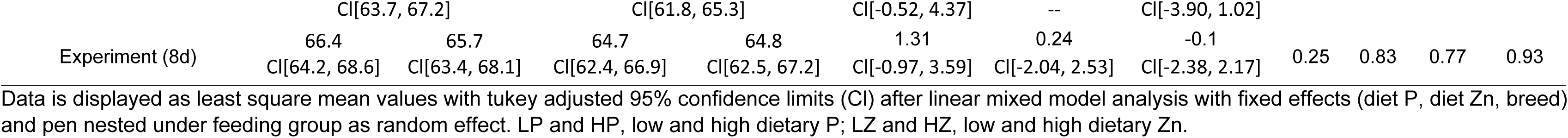
Response of cumulative feed intake and laying performance of adult laying hens to varying dietary phosphorus supply (0.37% or 0.84%) during two weeks pre-feeding and additional variation in dietary zinc supply (28 or 131 mg/kg) during subsequent eight days of experimental feeding.

Variation in diet P during acclimatization as well as diet P and diet Zn during the experimental phase did not affect feed intake of live weight of hens (P > 0.5 in any case). Average live weight of white vs. brown hens ranging between 1676-1959 g/hen vs. 1721-2811 g/hen as well as 1652-1943 g/hen vs. 1776 vs. 2804 g/hen during acclimatization as well as experimental phase, respectively. The average cumulative feed intake ranged between 1696-3026 g/pen and 1890-2839 g/pen per week of acclimatization and 8 d of experimental phase, respectively. The only relevant effect could be estimated for the breed of hen, with white hens showing on average - 313 Cl[-411; -215] g and -290 Cl[-392; -188] g lower live weight than brown hens during acclimatization and experimental phase, respectively (P < 0.0001). This presumably also affected the feed intake accordingly, but the present setup did not allow for dissection of genetic effects on feed intake behavior.

Variatons in P feeding during acclimatization did not affect cumulative egg counts over 7 d with LSmeans of 6.88 Cl[6.73, 7.04] eggs/hen and 6.80 Cl[6.64, 6.95] eggs/hen, accounting for 98% and 97% laying performance of LP and HP hens (P = 0.43 in any case). The same was observed for the combined variation of P and Zn during the acclimatization phase with respective group LSmeans of 7.67, 8.01, 7.67, and 7.83 accounting for 96%, 100%, 96%, and 98% of LPLZ, LPHZ, HPLZ, and HPHZ, respectively. Numerically lower laying performance was observed in LZ hens of -0.25 Cl[-0.55, 0.05] eggs/hen over 8 d, which was not statistically confirmed (P_P_ = 0.55 and P_Zn_ = 0.10). In addition. laying performance did not differ between breeds during acclimatization nor experimental phase (P = 0.81 and 0.27, respectively). Egg weight was numerically reduced in HP hens (63.5 Cl[61.8, 65.3] g/egg) compared to LP hens (65.4 Cl[63.7, 67.2] g/egg) throughout the acclimatization phase, which was not statistically confirmed (P = 0.12). Neither diet P nor Zn affected the egg weights during the experimental phase which were 66.4 Cl[64.2, 68.6], 65.7 Cl[63.4, 68.1], 64.7 Cl[62.4, 66.9], and 64.8 Cl[62.5, 67.2] g/egg for LPLZ, LPHZ, HPLZ, and HPHZ hens, respectively (P_P_ = 0.25, P_Zn_ = 0.83). Breed did non affect egg weight significantly at any time point of the study period (P > 0.14 in all cases).

### Endogenous mineral status

Table 4 outlines the effects of dietary P and Zn on the endogenous calcium status in the end of the experimental phase as assessed via Ca concentrations in blood plasma, liver, femur, and eggs. Blood plasma Ca levels remained consistent across dietary treatments at 0.30-0.32 mg/mL, showing no significant variations due to dietary P, Zn or breed (P_P_ = 0.25, P_Zn_ = 0.45, P_P*Zn_ = 0.77, P_Breed_ = 0.08).

**Table 4.**
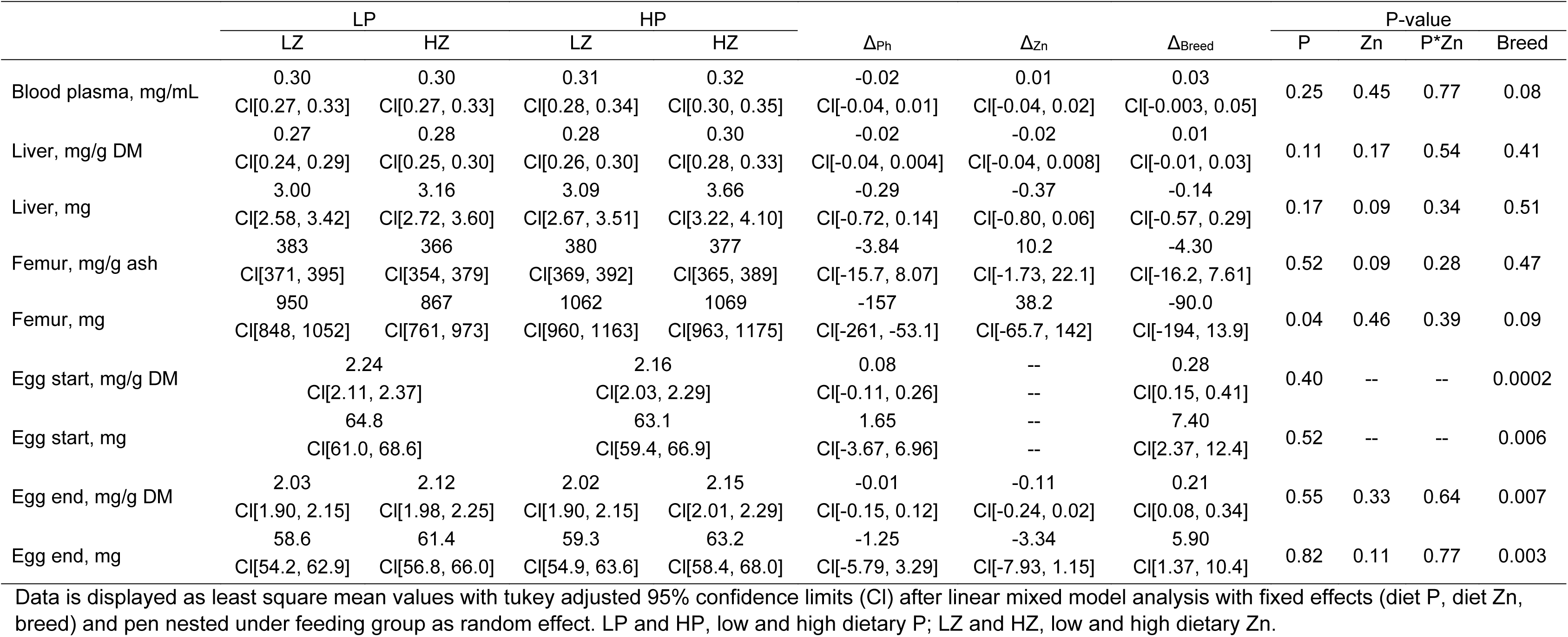
Response of endogenous calcium status in blood plasma, liver, and femur as well as its excretion with whole egg of laying hens at the beginning and end of the experiment after 14 d of varying dietary phosphorus (0.37% or 0.84%) prefeeding and 8d of additional variation in diet zinc supply (28 or 131 mg/kg).

While Ca liver concentrations did not differ between treatment groups or breeds (0.27-0.30 mg/g DM; P_P_ = 0.11, P_Zn_ = 0.17, P_P*Zn_ = 0.54, P_Breed_ = 0.41), absolute Ca content in the liver numerically decreased by -0.37 Cl[-0.80; 0.06] mg in LZ hens (P_Zn_ = 0.09) with no effect of or interaction with P (P= or breed.

Calcium concentration in the femur ash numerically increased by 10.2 Cl[-1.73; 22.1] mg/g ash in LZ hens (P_Zn_ = 0.09, P_P_ = 0.52, P_P*Zn_ = 0.28, P_Breed_ = 0.47) while the absolute femur Ca content significantly decreased by -157 Cl[-261; -53.1] mg with decreasing P in the diet (P = 0.04) and irrespective of the diet Zn supply or breed (P_Zn_ = 0.46, P_P*Zn_ = 0.39, P_Breed_ = 0.09). Hens started with comparable concentrations and contents of Ca in whole egg into the experiment, given by the respective data obtained in the beginning of the experimental phase in LP (2.24 Cl[2.11, 2.37] mg/g DM, 64.8 Cl[61.0, 68.6] mg) and HP hens (PP = 0.40 and 0.52). At the end of the experiments hens of the different feeding groups showed changes in egg Ca measures associated to P feeding (P = 0.55 and 0.82 for concentration and content, respectively) and just a numerical differences with respect to Zn feeding (P = 0.33 and 0.11 for concentration and content, respectively) with HZ hens having higher absolute contents of Ca in eggs (concentration: 2.03 Cl[1.90, 2.15], 2.12 Cl[1.98, 2.25], 2.02 Cl[1.90, 2.15], 2.15 Cl[2.01, 2.29]; content: 58.6 Cl[54.2, 62.9], 61.4 Cl[56.8, 66.0], 59.3 Cl[54.9, 63.6], 63.2 Cl[58.4, 68.0], in LPLZ, LPHZ, HPLZ, HPHZ hens, respectively.). The only factor affecting Ca in whole egg throughout the experimental phase was breed, with white hens showing higher concentrations (Δ_Breed_: 0.28 Cl[0.15, 0.41] and 0.21 [0.08, 0.34] mg/g DM at start and end of the experiment) and contents (Δ_Breed_: 7.40 Cl[2.37, 12.4] and 5.90 Cl[1.37, 10.4] mg at start and end of the experiment) than brown hens (P = 0.0002, 0.006, 0.007, and 0.003 for concentration and content at experimental start and end, respectively).

Table 5 highlights the effects of dietary P and Zn on the endogenous P status in the end of the experimental phase as assessed via P concentrations in blood plasma, liver, femur, and eggs. Blood plasma and liver P concentrations remained stable across diet groups between 0.51-0.60 mg/mL and 11.8-12.5 mg/g DM, respectively, with no significant impact from dietary P (P = 0.35 and 0.81 for plasma and liver, respectively) or Zn (P = 0.20 and 0.26 for plasma and liver, respectively). Liver concentrations were increased by 0.57 Cl[0.05, 1.09] mg/g DM in white compared to brown hens (P = 0.03). However, absolute liver P contents decreased by -11.3 CI[-22.5, -0.04] mg in LZ compared to HZ hens (P = 0.05).

**Table 5.**
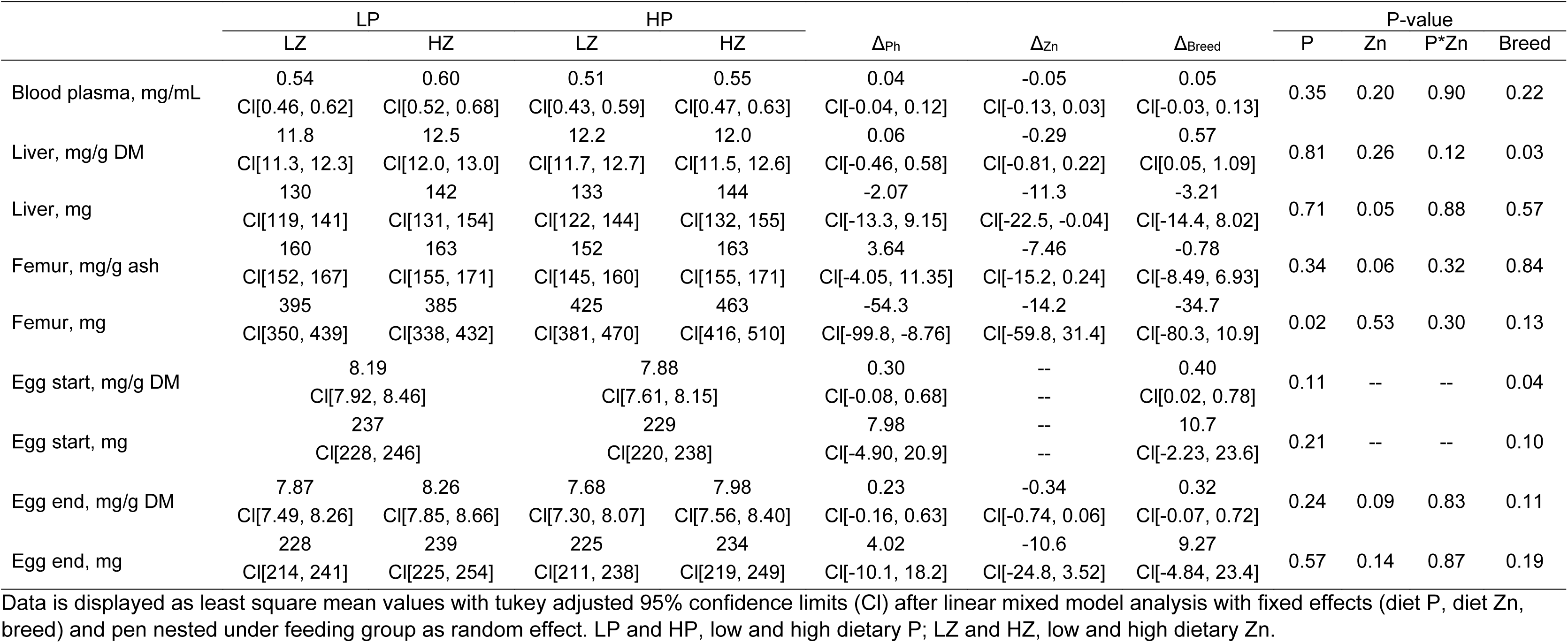
Response of endogenous phosphorus status in blood plasma, liver, and femur as well as its excretion with whole egg of laying hens at the beginning and end of the experiment after 14 d of varying dietary phosphorus (0.37% or 0.84%) prefeeding and 8d of additional variation in diet zinc supply (28 or 131 mg/kg).

| | LP | | HP | | $\Delta_{Ph}$ | $\Delta_{Zn}$ | $\Delta_{Breed}$ | P-value | | | |
| --- | --- | --- | --- | --- | --- | --- | --- | --- | --- | --- | --- |
|  | LZ | HZ | LZ | HZ |  |  |  | P | Zn | P*Zn | Breed |
| Blood plasma, mg/mL | 0.54<br>CI[0.46, 0.62] | 0.60<br>CI[0.52, 0.68] | 0.51<br>CI[0.43, 0.59] | 0.55<br>CI[0.47, 0.63] | 0.04<br>CI[-0.04, 0.12] | -0.05<br>CI[-0.13, 0.03] | 0.05<br>CI[-0.03, 0.13] | 0.35 | 0.20 | 0.90 | 0.22 |
| Liver, mg/g DM | 11.8<br>CI[11.3, 12.3] | 12.5<br>CI[12.0, 13.0] | 12.2<br>CI[11.7, 12.7] | 12.0<br>CI[11.5, 12.6] | 0.06<br>CI[-0.46, 0.58] | -0.29<br>CI[-0.81, 0.22] | 0.57<br>CI[0.05, 1.09] | 0.81 | 0.26 | 0.12 | 0.03 |
| Liver, mg | 130<br>CI[119, 141] | 142<br>CI[131, 154] | 133<br>CI[122, 144] | 144<br>CI[132, 155] | -2.07<br>CI[-13.3, 9.15] | -11.3<br>CI[-22.5, -0.04] | -3.21<br>CI[-14.4, 8.02] | 0.71 | 0.05 | 0.88 | 0.57 |
| Femur, mg/g ash | 160<br>CI[152, 167] | 163<br>CI[155, 171] | 152<br>CI[145, 160] | 163<br>CI[155, 171] | 3.64<br>CI[-4.05, 11.35] | -7.46<br>CI[-15.2, 0.24] | -0.78<br>CI[-8.49, 6.93] | 0.34 | 0.06 | 0.32 | 0.84 |
| Femur, mg | 395<br>CI[350, 439] | 385<br>CI[338, 432] | 425<br>CI[381, 470] | 463<br>CI[416, 510] | -54.3<br>CI[-99.8, -8.76] | -14.2<br>CI[-59.8, 31.4] | -34.7<br>CI[-80.3, 10.9] | 0.02 | 0.53 | 0.30 | 0.13 |
| Egg start, mg/g DM | 8.19<br>CI[7.92, 8.46] |  | 7.88<br>CI[7.61, 8.15] |  | 0.30<br>CI[-0.08, 0.68] | --<br>-- | 0.40<br>CI[0.02, 0.78] | 0.11 | -- | -- | 0.04 |
| Egg start, mg | 237<br>CI[228, 246] |  | 229<br>CI[220, 238] |  | 7.98<br>CI[-4.90, 20.9] | --<br>-- | 10.7<br>CI[-2.23, 23.6] | 0.21 | -- | -- | 0.10 |
| Egg end, mg/g DM | 7.87<br>CI[7.49, 8.26] | 8.26<br>CI[7.85, 8.66] | 7.68<br>CI[7.30, 8.07] | 7.98<br>CI[7.56, 8.40] | 0.23<br>CI[-0.16, 0.63] | -0.34<br>CI[-0.74, 0.06] | 0.32<br>CI[-0.07, 0.72] | 0.24 | 0.09 | 0.83 | 0.11 |
| Egg end, mg | 228<br>CI[214, 241] | 239<br>CI[225, 254] | 225<br>CI[211, 238] | 234<br>CI[219, 249] | 4.02<br>CI[-10.1, 18.2] | -10.6<br>CI[-24.8, 3.52] | 9.27<br>CI[-4.84, 23.4] | 0.57 | 0.14 | 0.87 | 0.19 |

Data is displayed as least square mean values with tukey adjusted 95% confidence limits (CI) after linear mixed model analysis with fixed effects (diet P, diet Zn, breed) and pen nested under feeding group as random effect. LP and HP, low and high dietary P; LZ and HZ, low and high dietary Zn.

Femur P concentration was not affected by diet P (P = 0.34) but reduced in LZ compared to HZ hens at -7.46 Cl[-15.2, 0.24] mg/kg ash (P = 0.06). Femur P content on the other hand was reduced in LP compared to HP hens by -54.3 Cl[-99.8, -8.76] mg (P = 0.02).

Comparable to the egg Ca data described above, egg phosphorus levels at experimental start were comparable between groups (7.88-8.19 mg/g DM and 229-237 mg for P concentration and content) (P = 0.11 and 0.21 for concentration and content). At the end of the experiment no significant effect of the P feeding could be observed on both P concentration and content in whole egg (P = 0.24 and 0.57). However, the Zn feeding regime appeared to shift P distribution to some extend with LZ hens an increase in concentration and content of -0.34 Cl[-0.74, 0.06] mt/g DM and -10.6 Cl[-24.8, 3.52] mg, respectively (P = 0.09 and 0.14, respectively). Although white hens again showed higher concentrations and contents of P in eggs at start and end of the experiments, this could only be statistically confirmed for the concentration at experimental start where white hens exceeded the level of brown hens by 0.40 Cl[0.02, 0.78] mg/g DM (P = 0.04).

The effects of dietary P and Zn on the endogenous Zn status in the end of the experimental phase as assessed via Zn concentrations in blood plasma, liver, femur, and eggs are shown in Table 6. The LZ compared to HZ hens had decreased plasma Zn concentrations by -0.83 CI[-1.29; -0.36] mg/L (P = 0.0008). Otherwise, no significant effects of P, breed, and interaction was observed (P_P_ = 0.19, P_P*Zn_ = 0.28, P_Breed_ = 0.23).

**Table 6.** Response of endogenous zinc status in blood plasma, liver, and femur as well as its excretion with whole egg of laying hens at the beginning and end of the experiment after 14 d of varying dietary phosphorus (0.37% or 0.84%) prefeeding and 8d of additional variation in diet zinc supply (28 or 131 mg/kg).

| | LP | | HP | | $\Delta_{Ph}$ | $\Delta_{Zn}$ | $\Delta_{Breed}$ | P-value | | | |
| --- | --- | --- | --- | --- | --- | --- | --- | --- | --- | --- | --- |
|  | LZ | HZ | LZ | HZ |  |  |  | P | Zn | P*Zn | Breed |
| Blood plasma, mg/L | 4.59<br>CI[4.14, 5.04] | 5.17<br>CI[4.69, 5.64] | 4.03<br>CI[3.58, 4.49] | 5.11<br>CI[4.64, 5.58] | 0.31<br>CI[-0.16, 0.77] | -0.83<br>CI[-1.29, -0.36] | 0.28<br>CI[-0.18, 0.74] | 0.19 | 0.0008 | 0.28 | 0.23 |
| Liver, mg/kg DM | 93.6<br>CI[88.4, 98.9] | 102<br>CI[96.3, 107] | 95.2<br>CI[90.0, 100] | 101<br>CI[95.1, 106] | -0.19<br>CI[-5.56, 5.16] | -6.78<br>CI[-12.1, -1.42] | 3.77<br>CI[-1.59, 9.14] | 0.94 | 0.01 | 0.60 | 0.16 |
| Liver, mg | 1.04<br>CI[0.94, 1.13] | 1.16<br>CI[1.06, 1.26] | 1.04<br>CI[0.94, 1.14] | 1.19<br>CI[1.09, 1.29] | -0.02<br>CI[-0.12, 0.08] | -0.14<br>CI[-0.23, -0.04] | -0.03<br>CI[-0.13, 0.06] | 0.71 | 0.008 | 0.77 | 0.50 |
| Femur, mg/kg ash | 490<br>CI[401, 579] | 535<br>CI[442, 628] | 585<br>CI[496, 674] | 606<br>CI[513, 699] | -82.4<br>CI[-173, 8.54] | -32.9<br>CI[-124, 58.2] | -57.9<br>CI[-149, 33.1] | 0.07 | 0.47 | 0.79 | 0.21 |
| Femur, mg | 1.21<br>CI[0.96, 1.47] | 1.23<br>CI[0.96, 1.50] | 1.64<br>CI[1.38, 1.89] | 1.73<br>CI[1.47, 2.00] | -0.46<br>CI[-0.72, -0.20] | -0.06<br>CI[-0.32, 0.20] | -0.28<br>CI[-0.54, -0.02] | 0.0009 | 0.66 | 0.77 | 0.03 |
| Egg start, mg/kg DM | 55.5<br>CI[52.4, 58.6] |  | 51.1<br>CI[48.0, 54.2] |  | 4.42<br>CI[0.04, 8.80] | -- | 4.85<br>CI[1.21, 8.48] | 0.05 | -- | -- | 0.01 |
| Egg start, mg | 1.61<br>CI[1.52, 1.70] |  | 1.49<br>CI[1.39, 1.58] |  | 0.12<br>CI[-0.008, 0.25] | -- | 0.06<br>CI[0.01, 0.25] | 0.06 | -- | -- | 0.03 |
| Egg end, mg/kg DM | 48.6<br>CI[44.8, 52.3] | 54.4<br>CI[50.4, 58.3] | 46.9<br>CI[43.2, 50.7] | 51.4<br>CI[47.3, 55.5] | 2.30<br>CI[-1.59, 6.19] | -5.14<br>CI[-9.03, -1.24] | 3.46<br>CI[-0.43, 7.34] | 0.24 | 0.01 | 0.74 | 0.08 |
| Egg end, mg | 1.40<br>CI[1.29, 1.52] | 1.58<br>CI[1.46, 1.70] | 1.37<br>CI[1.25, 1.48] | 1.51<br>CI[1.38, 1.63] | 0.05<br>CI[-0.07, 0.17] | -0.16<br>CI[-0.27, -0.04] | 0.10<br>CI[-0.02, 0.22] | 0.39 | 0.01 | 0.77 | 0.10 |

Data is displayed as least square mean values with tukey adjusted 95% confidence limits (CI) after linear mixed model analysis with fixed effects (diet P, diet Zn, breed) and pen nested under feeding group as random effect. LP and HP, low and high dietary P; LZ and HZ, low and high dietary Zn.

Liver Zn concentrations decreased in LZ hens by -6.78 mg/kg DM CI[ -12.1, -1.42] (P = 0.01) compared to high Zn diets, without any effects from P, breed, or interaction (P_P_ = 0.94, P_P*Zn_ = 0.60, P_Breed_ = 0.16). This response pattern was comparable for the liver Zn content but more pronounced with LZ hens showing a drop of -0.14 Cl[-0.23, -0.04] mg compared to HZ hens (P = 0.0008) and no effect of P, breed or interaction (P_P_ = 0.71, P_P*Zn_ = 0.77, P_Breed_ = 0.50). Femur Zn concentrations were reduced by -82.4 Cl[-173, 8.54] mg/kg ash in LP compared to HP hens (P = 0.07) but were affected by diet Zn (P = 0.47), breed (P = 0.21) or interaction (P = 0.79). This effect was much more pronounced when looking at the Zn content in the femur which was -0.46 Cl[-0.72, -0.20] mg lower in LP compared to HP hens (P = 0.0009) without effects of diet Zn (P = 0.66) and interaction (P = 0.77) but a significantly reduced levels in white compared to brown hens by -0.28 Cl[-0.54, -0.02] mg (P = 0.03).

The varying P feeding during acclimatization affected both the concentration and content of Zn in whole egg at the start of the experiment, with LP hens showing 4.42 Cl[0.04, 8.80] mg/kg DM and 0.12 [-0.008, 0.25] mg higher levels than HP hens (P = 0.05 and 0.06 for concentration and content). In addition, white hens lost more Zn with whole egg than brown hens at the start of the experiment as indicated by an increase in concentration and content of 4.85 Cl[1.21, 8.48] mg/kg DM and 0.06 Cl[0.01, 0.25] mg (P = 0.01 and 0.03 for concentration and content). After 8 d of experimental feeding varying levels of P and Zn, whole egg concentrations and contents of Zn significantly decreased with reduced diet Zn supply by -5.14 Cl[-9.03, -1.24] mg/kg DM and -0.16 Cl[-0.27, -0.04] mg (P = 0.01 in both cases) whereas P, breed, and interaction exerted no significant effects (P_P_ = 0.24 and 0.39, P_P*Zn_ = 0.74 and 0.77, P_Breed_ = 0.08 and 0.10).

### Quality of egg shells and bones

Table 7 highlights quality parameters of egg shells and femoral bones during 14 days of acclimatization feeding and 8 d of experimental feeding. Neither the thickness nor breaking strength of the egg shells were significantly affected by the feeding during the study period (P > 0.54 in any case). The same was true for the effect of breed on the thickness of egg shells, but not the egg breaking strength with white hens laying showing on average 1.29 Cl[0.87, 1.71] kg and 1.28 Cl[0.84, 1.72] kg higher breaking strength during acclimatization and experiment than brown hens, respectively (P < 0.0001 in any case).

**Table 7.** Response of egg shell and bone quality of adult laying hens to varying dietary phosphorus supply (0.37% or 0.84%) during two weeks Pre-feeding and additional variation in dietary zinc supply (28 or 131 mg/kg) during eight days of experimental feeding.

| | LP | | HP | | $\Delta_{Ph}$ | $\Delta_{Zn}$ | $\Delta_{Breed}$ | P | | | |
| --- | --- | --- | --- | --- | --- | --- | --- | --- | --- | --- | --- |
|  | LZ | HZ | LZ | HZ |  |  |  | Ph | Zn | Ph*Zn | Breed |
| Egg shell thickn., mm |  |  |  |  |  |  |  |  |  |  |  |
| Pre-feeding |  |  |  |  |  |  |  |  |  |  |  |
| Week 1 | 0.40 |  | 0.41 |  | -0.001 | -- | -0.003 | 0.87 | -- | -- | 0.60 |
|  | CI[0.39, 0.41] |  | CI[0.39, 0.42] |  | CI[-0.02, 0.01] | -- | CI[-0.02, 0.01] |  |  |  |  |
| Week 2 | 0.40 |  | 0.40 |  | 0.001 | -- | 0.006 | 0.83 | -- | -- | 0.34 |
|  | CI[0.40, 0.41] |  | CI[0.39, 0.41] |  | CI[-0.01, 0.01] | -- | CI[-0.006, 0.02] |  |  |  |  |
| Total | 0.41 |  | 0.40 |  | 0.002 | 0.0009 |  | 0.79 | -- | -- | 0.23 |
|  | CI[0.40, 0.41] |  | CI[0.39, 0.41] |  | CI[-0.01, 0.01] | CI[-0.01, 0.01] |  |  |  |  |  |
| Experiment | 0.41 | 0.40 | 0.41 | 0.41 | -0.003 | 0.003 | 0.002 | 0.57 | 0.54 | 0.57 | 0.75 |
|  | CI[0.40, 0.42] | CI[0.39, 0.41] | CI[0.40, 0.42] | CI[0.40, 0.42] | CI[-0.01, 0.008] | CI[-0.008, 0.01] | CI[-0.009, 0.01] |  |  |  |  |
| Egg shell strength, kg |  |  |  |  |  |  |  |  |  |  |  |
| Pre-feeding |  |  |  |  |  |  |  |  |  |  |  |
| Week 1 | 6.11 |  | 6.23 |  | -0.11 | -- | 1.22 | 0.63 | -- | -- | <0.0001 |
|  | CI[5.77, 6.46] |  | CI[5.88, 6.57] |  | CI[-0.60, 0.37] | -- | [0.73, 1.71] |  |  |  |  |
| Week 2 | 5.79 |  | 5.90 |  | -0.12 | -- | 1.35 | 0.64 | -- | -- | <0.0001 |
|  | CI[5.41, 6.16] |  | CI[5.53, 6.28] |  | CI[-0.64, 0.41] | -- | CI[0.89, 1.81] |  |  |  |  |
| Total | 5.95 |  | 6.06 |  | -0.11 | -- | 1.29 | 0.63 | -- | -- | <0.0001 |
|  | CI[5.61, 6.29] |  | CI[5.72, 6.40] |  | CI[-0.59, 0.37] | -- | CI[0.87, 1.71] |  |  |  |  |
| Experiment | 5.99 | 5.73 | 5.77 | 5.84 | 0.05 | 0.09 | 1.28 | 0.81 | 0.67 | 0.44 | <0.0001 |
|  | CI[5.56, 6.42] | CI[5.28, 6.18] | CI[5.34, 6.20] | CI[5.40, 6.30] | CI[-0.39, 0.49] | CI[-0.35, 0.53] | CI[0.84, 1.72] |  |  |  |  |
| Femur |  |  |  |  |  |  |  |  |  |  |  |
| Length, cm | 8.16 | 8.01 | 7.99 | 8.17 | -0.0008 | -0.01 | -0.53 | 0.99 | 0.82 | 0.02 | <0.0001 |
|  | CI[8.03, 8.29] | CI[7.87, 8.14] | CI[7.86, 8.12] | CI[8.04, 8.31] | CI[-0.14, 0.13] | CI[-0.15, 0.12] | CI[-0.66, -0.39] |  |  |  |  |
| Diameter, cm | 0.76 | 0.74 | 0.78 | 0.76 | -0.02 | 0.02 | -0.10 | 0.19 | 0.32 | 0.95 | <0.0001 |
|  | CI[0.73, 0.79] | CI[0.71, 0.77] | CI[0.75, 0.81] | CI[0.73, 0.80] | CI[-0.05, 0.01] | CI[-0.02, 0.05] | CI[-0.13, -0.07] |  |  |  |  |
| Strength, kg | 215 | 215 | 216 | 242 | -14.2 | -12.8 | 17.6 | 0.37 | 0.41 | 0.39 | 0.26 |
|  | CI[185, 246] | CI[183, 247] | CI[186, 247] | CI[210, 274] | CI[-45.4, 17.1] | CI[-44.1, 18.5] | CI[-13.7, 48.9] |  |  |  |  |
| Ash, g | 2.48 | 2.36 | 2.79 | 2.84 | -0.39 | 0.03 | -0.21 | 0.006 | 0.80 | 0.54 | 0.13 |
|  | CI[2.22, 2.75] | CI[2.09, 2.64] | CI[2.53, 3.07] | CI[2.56, 3.11] | CI[-0.66, -0.12] | CI[-0.24, 0.31] | CI[-0.48, 0.06] |  |  |  |  |
| Ca:P, molar ratio | 1.88 | 1.74 | 1.94 | 1.79 | -0.05 | 0.15 | -0.006 | 0.33 | 0.01 | 0.90 | 0.91 |
|  | CI[1.77, 1.99] | CI[1.62, 1.85] | CI[1.83, 2.05] | CI[1.67, 1.90] | CI[-0.17, 0.06] | CI[0.04, 0.26] | CI[-0.12, 0.11] |  |  |  |  |

Data is displayed as least square mean values with tukey adjusted 95% confidence limits (CI) after linear mixed model analysis with fixed effects (diet P, diet Zn, breed) and pen nested under feeding group as random effect. LP and HP, low and high dietary P; LZ and HZ, low and high dietary Zn.thickn., thickness

The length, diameter, and strength of femoral bones was not affected by the P and Zn feeding individually (P_P_ = 0.99, 0.19, and 0.37; P_Zn_ = 0.82, 0.32, and 0.41 for length, diameter, and strength, respectively) but a significant interaction between P and Zn feeding (P = 0.02) was observed for bone length due to inverse ratios of bone length of LZ:HZ hens in the LP and HP cluster, respectively (ratios were 1.01 and 0.98 for LZ:HZ in LP and HP, respectively). Both length and diameter but not breaking strength were different between breeds (P < 0.0001, <0.0001, = 0.26, respectively), with white hens having -0.53 Cl[-0.66, -0.39] cm and -0.10 Cl[-0.13, -0.07] cm shorter and narrower bones, respectively.

Total bone ash content differed with respect to the P feeding at the end of the experiment with LP hens having -0.39 Cl[-0.66, -0.12] g less total ash in their femoral bones than HP hens (P = 0.006). At the same time, diet Zn, breed, and interaction showed no effects (P_Zn_ = 0.80, P_Breed_ = 0.13, P_P*Zn_ = 0.54).

The femoral Ca:P molar ratio at the end of the experiment differed in response to the Zn feeding regime, where LZ hens showed a wider ratio by 0.15 Cl[0.04, 0.26] compared to HZ hens (P = 0.01) while diet P, breed, and interaction had no effect (P_P_ = 0.33, P_Breed_ = 0.91, P_P*Zn_ = 0.90).

## Discussion

Earlier, we developed (Brugger et al., 2014) and validated (Boerboom et al., 2022) an experimental model of subclinical Zn deficiency in weaned piglets involving 14 d acclimatization feeding according to recommendations followed by 8 d of varying Zn supply to a phytate-rich diet. Such models are crucial to establish Zn requirements and study metabolic principles without pathological bias (Brugger et al., 2022). For chickens, such a model has yet to be developed. Due to their higher demand for calcium (Lohmann, 2020a, 2020b; Aviagen, 2022), layers differ in metabolism and feeding conditions from broilers. Since broilers showed a relevant capacity for intestinal phytate degradation in earlier studies (Zeller et al., 2015; Sommerfeld et al., 2019) a higher resilience to diet-induced Zn deficiency can be assumed as compared to weaned piglets (Schlegel et al., 2010, 2013; Boerboom, 2021). Although layers generally have lower phytate-P utilization than broilers they follow the same trends in response to varying mineral phosphorus supply with high mineral P supply depressing phytate degradation (Ma et al., 2019; Sommerfeld et al., 2020). The present study attempted to test the susceptibility of adult laying hens to dietary shortage in mineral Zn after a high versus low mineral P acclimatization feeding. Since earlier studies further suggested differences in phytate breakdown of different layer strains, we included Lohmann Brown Classic and Lohmann LSL Classic hens in the study.

### Response to reduced diet Zn irrespective of mineral P supply

By the end of the present study, plasma, liver, and egg Zn levels significantly decreased with lower Zn intake. Plasma Zn can be affected by multiple factors other than Zn, such as inflammation or reduced feed intake (Suttle, 2022). However, an inflammation-dependent decrease in circulatory Zn would have probably been accompanied by a rise in liver Zn as suggested by studies in rodents and in 4-wk-old white Leghorn chicks (Klasing, 1984; Milanino et al., 1986; Cousins and Leinart, 1988; Shea-Budgell et al., 2006). In our study, both plasma, liver and egg Zn decreased, with no observed effects on feed intake or pathologies. After gut absorption, Zn is transported via the portal vein to the liver for mid-term storage or immediate distribution (Suttle, 2022). Previous studies suggested that liver Zn concentration can reflect Zn supply in piglets as liver Zn only accumulates in case of sufficient and surplus Zn supply, whereas insufficient Zn intake causes a plateau at minimum level (Brugger et al., 2014; Boerboom et al., 2022). However, no further increase was observed when adding phytase while increasing zinc supplementation from 30 to 90 mg/kg diet in broilers (Philippi et al., 2023a). Similar data for laying hens is currently not available. Paulicks and Kirchgessner (1994) demonstrated the homeostatic regulation of the laying hen limits the accumulation of Zn in eggs when receiving diet Zn above requirements, and otherwise depletion of egg Zn was observed when the Zn supply was reduced below the requirement threshold for a sufficient amount of time (Paulicks and Kirchgessner, 1994). Our study found similar plasma and liver Zn responses as described for piglets (Brugger et al., 2014; Boerboom et al., 2022). Comparably to these studies in piglets, plasma and liver Zn, but also egg Zn levels, decreased without affecting performance, indicating a subclinical Zn deficiency-state after only 8d of reduced alimentary supply. In contrast, earlier data in laying hens suggest that about 4 months of a Zn-deficient diet are needed to induce clinical signs of Zn deficiency such as reduced egg production and feed intake (Paulicks and Kirchgessner, 1994). However, this data was obtained in layers that do not represent nowadays genotypes and it can be assumed that the susceptibility of modern breeds to dietary Zn deficiency increased with their performance. Overall, our laying hens appeared less susceptible to dietary Zn deficiency than pigs, as indicated for example by the femur Zn depletion (5.7% compared to 24.7% in piglets after 8 d of challenge) which numerically decreased in the present study but could not be statistically confirmed (Brugger et al., 2014). However, the observed depletion rate of 5.7% as affected by diet Zn variation might be biased by the fact that we used the complete femur instead of the just the femoral head as in our pig studies, which according to earlier studies e.g. in rats (Windisch et al., 2002) contains the majority of the highly mobile bone zinc pool. The total extent of diet Zn-dependent bone Zn depletion in the layer might therefore be more pronounced than the present study suggests (please see a more thorough discussion of the matter below in our subsection on the limitations of our study). Nevertheless, the earlier reported time frame of 10-12 d until weaned piglets develop a clinical Zn deficiency is probably longer in the laying hen, assuming a comparable percentage of mobilizable Zn in their skeleton as compared to rats and pigs (Windisch and Kirchgessner, 1994; Boerboom et al., 2022).

In summary, the present study demonstrated that an 8-day feeding of only native Zn induced subclinical Zn deficiency in laying hens as evidenced by the reduction in bone, and especially liver and egg Zn while no clinical symptoms of Zn deficiency were observed. This indicates a greater susceptibility to Zn malnutrition than broilers, likely due to lower digestive capacity for phytate and constantly high endogenous Zn loses via eggs as discussed above.

### Consequences of reduced mineral P supply for diet Zn utilization

A major debate in the poultry science community is how well chickens can use phytate P. Corn and soybean meal were the major diet ingredients in our study and have high phytate P concentration (De Boland et al., 1975; Humer et al., 2015). Broiler chickens generally show high levels of phytate disappearance, indicating that endogenous mucosal and bacterial phytases might play an important role in phytate hydrolysis (Rodehutscord et al., 2017). Strong evidence to support this claim comes from targeted metabolic studies were phytate disappearance alongside the gastrointestinal tract of wildtype and gnotobiotic ROSS broilers has been investigated, suggesting an overall high endogenous potential to utilize phytate-P even in the absence of a gastrointestinal microbiome, which is impaired by the addition of mineral P to the diet (Zeller et al., 2015; Sommerfeld et al., 2019). Generally, phytate degradation in broilers is overall higher than in laying hens which may be attributed to differences in retention time of digesta and a higher Ca concentration in the layer’s digestive tract (Ma et al., 2019). The higher Ca content in layer diets may impair phytate degradation by increasing the intestinal pH and the likelihood for the formation of phytase-resistant Ca-Zn-phytates (Humer et al., 2015; Ma et al., 2019; Philippi et al., 2023b), thereby reducing the Zn and P availability (Philippi et al., 2023b).

Liver Zn responds to the diet Zn supply status (Brugger et al., 2022), yet no significant variation in liver Zn with varying P was observed, which is in strong contrast to our initial hypothesis that reduced P feeding may increase Zn uptake due to an improved phytate breakdown in the gastrointestinal tract. Diet Zn on the other hand did affect liver Zn with significantly reduced values after 8 d feeding without mineral Zn supplementation irrespective of the diet P supply. Since we did not staged slaughtering of hens throughout the study period, it is unclear if the P feeding affected diet Zn utilization in the beginning of the acclimatization feeding. Should reduced mineral P feeding have increased the levels of absorbable Zn over time, this probably would have redirected surplus uptake to the excretion into the gastrointestinal tract and insufficient supply would have been compensated by more efficient absorption (Brugger and Windisch, 2019). This stresses the need for follow up studies mapping the Zn absorption and liver Zn status over time. In addition, our observation of mineral P impairing the bone Zn concentration and content irrespective of the diet Zn supply may have increased serum Zn, thereby misleading the system into reducing gut Zn absorption, a hypothesis needing further investigation.

### Interactions of mineral P and Zn with respect to bone and mineral metabolism

Reduced P intake decreased femur Zn, Ca and P contents and concentrations, though this effect was not seen when related to the concentration in femur ash, likely due to bone resorption reducing the overall mineral content of the skeleton. The reduced alimentary P supply seemed to promote bone resorption, indicated by an overall decrease in bone ash content in our study. Tibia zinc levels in compact bone and bone marrow of the tibia in laying hens declined in low P diets without phytase (Martinez Rojas et al., 2018). Despite increased bone resorption, our study showed no improvement in Zn utilization. However, low Zn feeding tended to reduce P and increase Ca in femur ash, raising the Ca-P molar ratio. The reduced P content in bone suggests a potential impact of Zn on the organic bone matrix, which contains various phosphorylated proteins (Boskey, 2007). These effects were however less strong than the effects of low P feeding on femur minerals themselves. This suggests that P and Zn may affect bone mineral status independently, which would agree to earlier studies in a rat model of osteoporosis (Windisch et al., 2002). While it is well known, that Zn plays a role in bone metabolism by stimulating bone formation via alkaline phosphatase activity, collagen synthesis and inhibiting osteoclastic bone resorption (Yamaguchi, 1998), direct effects of Zn on P metabolism have scarcely been investigated.

At first glance, it may seem surprising that the reduction in bone mineralization did not significantly impact bone stability, as measured by bone-breaking strength tests. While some numerically lower values were observed for LP hens compared to HP hens, these differences were not statistically significant. This phenomenon can be explained by the unique physiological adaptations of laying hens and egg-laying birds in general. They develop a labile reservoir of calcium (Ca) and phosphorus (P) in the form of medullary bone tissue within the cavities of their long bones (Whitehead, 2004). Unlike structurally critical cortical and trabecular bone, medullary bone serves as a non-structural matrix that supports the high mineral demand for eggshell formation. This adaptation minimizes the need to draw minerals from structural bone tissue (Bonucci and Gherardi, 1975; Dacke et al., 1993). While medullary bone can enhance overall bone stability by bolstering the fracture resistance of surrounding cortical bone when present in large amounts (Fleming et al., 1998), its depletion does not immediately result in a drastic impairment of the structural stability of bone tissue. Only prolonged and severe dietary Ca or P deficiencies, typically lasting over two weeks, can fully deplete medullary bone reserves and force mineral mobilization from structural bone, as demonstrated in studies with quails (Dacke et al., 1993). Based on this, we conclude that the observed bone resorption primarily reflects changes in medullary bone tissue, resulting in numerical differences in bone-breaking strength without reaching statistical significance.

### Breed effects

We observed significant effects of breed on body weight, while egg yield and weight remained unaffected by breed. However, shell strength was increased in white laying hens, which correlated to a higher calcium content in the whole eggs in white laying hens. Moreover, egg Zn was increased in white laying hens compared to brown layers at the start of the experiment. We observed lower absolute femur Ca and Zn contents. This aligns with findings by Kraus et al. (2022), who reported variations in bone Zn content across different chicken breeds, with higher Zn levels in the femur than in the tibia (Kraus et al., 2022). Most likely constant efforts of the industry in selecting birds with optimal performance and health characteristics as observed in broiler chickens (Zuidhof et al., 2014), potentially had similar effects in laying hens, the extent of which is unclear at the moment. Moreover, changes in absolute femoral mineral contents may be explained by the reduced diameter and length of femur bones in white laying hens.

### Limitations of the study

Our study had several limitations. For animal welfare reasons and to exclude effects of single cages on study outcome, 2 hens (1 per breed) were housed together per cage. Therefore, it was not possible to determine the feed intake per animal. Moreover, several parameters could only be determined as an endpoint measurement in the end of the trial. In addition, the stabling system did not allow for a quantitative collection of the faecal matter, therefore apparent retention of nutrients and elements could not be assessed and due to the combined excretion of faecal and renal excrements in birds, an indicator-dilution approach was not possible. However, this study was conducted as a pilot to guide future research on the matter. These follow-up projects should target several time-points by including more hens and slaughter them serially. In addition, balance studies in metabolism cages should be conducted.

Our previous subclinical Zn-deficiency studies in piglets (Brugger et al., 2014) used the femoral head, containing high amounts of trabecular bone, which has been suggested in previous rat studies to contain the highly mobile Zn pool of bone tissue (Windisch et al., 2002). This was confirmed by recent yet unpublished studies in chickens applying nanoscale secondary ion mass spectrometry (nano-SIMS) (personal communication by Prof. Dirk Schaumlöffel, Université de Pau et des Pays de l’Adour / CNRS France). Therefore, our choice to ash the complete femur instead of just the femoral head may have diluted the matrix and potentially impaired the resolution of Zn exchange responses to diet Zn variation in addition to the potential overshadowing by the strong P effect on bone Zn. The notion behind ashing the complete femur was to increase the individual amount of bone ash for mineral analysis. In addition, as mentioned above laying hens develop medullary bone tissue primarily in the cavities of the long bones with are usually filled with air in birds to serves as an additional labile pool of Ca and P to support egg shell formation (Whitehead, 2004). Since we were also interested in observing the effects of our feeding regime on Ca and P exchange in bone, considering the complete bone for our analysis made sense then. Own and others’ follow-up attempts to reproduce and further understand the observations made in the present study, must therefore include a separate ashing of trabecular and medullar bone tissue of layers.

### Conclusion and outlook

Our study demonstrated that feeding a Zn-deficient diet for 8 days induced a subclinical zinc deficiency in laying hens, marked by reduced body Zn levels without promoting any clinical signs of Zn deficiency. This supports the applicability of our previously established pig model for inducing subclinical Zn deficiency in laying hens. However, the response within the 8 d of insufficient Zn feeding was milder than in piglets, suggesting a prolonged feeding model could be established to increase the resolution of measurements by strengthening the subclinical Zn deficiency stimulus. Follow-up studies should therefore focus on the adaption of Zn metabolism in the layer over time, to map the point where subclinical transforms to clinical zinc deficiency to better understand the kinetics. Contrary to our initial hypothesis, reducing mineral P did not improve Zn utilization under the present feeding conditions. Instead, reduced P decreased bone Zn status independently of dietary Zn, and increased the Ca-P molar ratio in bones, mainly by lowering femur ash P. To the best of our knowledge, these are novel interactions that have not been reported before, the precise functional background of which as well as consequences for animal health and productivity are yet unclear and warrant further research.

## Acknowledgements

We thank the technical personal and students of our institutes for excellent technical assistance, Phillip Urban M.Sc. (TUM).

## Funding information

No particular funding was obtained for the conduction of this study.

## Data availability statement

Raw data from this study can be obtained from the corresponding author upon request.

## Conflict of interest statement

Nothing to declare

## Author contribution

AL Writing-original draft, Visualization

AW Investigation

DB Writing-original draft, Supervision, Project Administration, Investigation, Conceptualization, Data curation, Formal analysis, Visualization

JP Investigation

JS Investigation, Supervision, Writing – Review and editing

KS Supervision, Resources, Writing – Review and editing

RP Investigation, Supervision, Writing – Review and editing

WW Resources, Writing – Review and editing

## Notes

### Competing Interest Statement

The authors have declared no competing interest.

