## Supplementary Table 1 for "Temporary suspension of mineral phosphorus reduces mobilizable bone zinc in adult laying hens irrespective of the dietary zinc supply"

Supplementary Table 1. Response of live weight of adult laying hens to varying dietary phosphorus supply (0.37% or 0.84%) during two weeks pre-feeding and additional variation in dietary zinc supply (28 or 131 mg/kg) during subsequent eight days of experimental feeding.

| | LP | | HP | | | | | P-value | | | |
|---|---|---|---|---|---|---|---|---|---|---|---|
| | LZ | HZ | LZ | HZ | $\Delta_{P}$ | $\Delta_{Zn}$ | $\Delta_{Breed}$ | P | Zn | P*Zn | Breed |
| Live weight | | | | | | | | | | | |
| kg/hen | | | | | | | | | | | |
| Pre-feeding | | | | | | | | | | | |
| Week 1 | 1972 | | 1971 | | 1.21 | -- | -326 | 0.98 | -- | -- | <0.0001 |
| | CI[1897, 2048] | | [1896, 2047] | | CI[-105, 108] | -- | CI[-424, -228] | | | | |
| Week 2 | 1939 | | 1930 | | 8.33 | -- | -300 | 0.87 | -- | -- | <0.0001 |
| | CI[1864, 2013] | | CI[1856, 2005] | | CI[-97.4, 114] | -- | CI[-400, -201] | | | | |
| Total (14 d) | 1955 | | 1951 | | 4.79 | -- | -313 | 0.93 | -- | -- | <0.0001 |
| | CI[1881, 2030] | | CI[1876, 2025] | | CI[-101, 110] | -- | CI[-411, -215] | | | | |
| Experiment (8 d) | 1965 | 1933 | 1904 | 1958 | 17.3 | -11.1 | -290 | 0.73 | 0.83 | 0.40 | <0.0001 |
| | CI[1863,2066] | CI[1831, 2035] | CI[1803, 2006] | CI[1857, 2060] | CI[-84.4, 119] | CI[-113, 90.6] | CI[-392, -188] | | | | |
